# Temporal Shifts in Antibiotic Resistance Elements Govern Virus-Pathogen Conflicts

**DOI:** 10.1101/2020.12.16.423150

**Authors:** Kristen N. LeGault, Stephanie G. Hays, Angus Angermeyer, Amelia C. McKitterick, Fatema-tuz Johura, Marzia Sultana, Munirul Alam, Kimberley D. Seed

**Affiliations:** Department of Plant and Microbial Biology, University of California, Berkeley, Berkeley, CA 94720, USA; icddr,b, formerly International Centre for Diarrhoeal Disease Research, Dhaka, Bangladesh; Chan Zuckerberg Biohub, San Francisco, CA 94158, USA

## Abstract

Bacteriophage predation has selected for widespread and diverse anti-phage systems that frequently cluster on mobilizable defense islands in bacterial genomes. Understanding phage-host co-evolutionary dynamics has lagged due to a dearth longitudinal sampling in natural environments. Here, using time shift experiments we show that epidemic *Vibrio cholerae* and lytic phages recovered from cholera patient stool samples display negative frequency-dependent co-evolution. We find that SXT integrative and conjugative elements (ICEs), which are infamous for conferring antibiotic resistance, invariably encode phage defense systems. SXT ICEs in *V. cholerae* govern susceptibility to phages in clinical samples and we demonstrate phage counter-adaptation to SXT ICE restriction over a three-year sampling period. Further, phage infection stimulates high frequency SXT ICE conjugation, leading to the concurrent dissemination of phage and antibiotic resistance.

## Main Text

Bacteriophage (phage) predation has been estimated to account for upwards of 40% of all bacterial mortality (1), shaping microbial community composition and selecting for bacteria that evade predation (2, 3). Co-evolution between phages and their hosts contributes to global biodiversity (2, 4–6). Indeed, genetic differences between closely related strains are largely phage resistance genes (7), with anti-phage defense systems comprising upwards of 10% of a bacterial genome (8), and mutations that decrease host susceptibility accounting for further inter-strain variability (9, 10). Phylogenetic analyses show that defense systems associate with mobilome genes, indicating that phage predation can drive bacterial evolution through mobile genetic element (MGE) flux (8, 11). A recent explosion in the identification of novel defense systems (11, 12) has allowed for a re-assessment of bacterial genomes, identifying resistance determinants in genetic contexts not previously associated with phage defense.

Despite the central role of phages in microbial evolution and ecology, there remains a dearth of biologically relevant systems to study ongoing phage-host interactions in natural environments. Approaches to study phage resistance systems often rely on ectopic expression of candidate systems in heterologous bacterial hosts and assessing restriction of phages that are not known to have encountered the system in nature. Although this approach has proven lucrative for the discovery of novel defense systems (11–14) and is especially valuable for assessing genes from unculturable organisms or those without known phages, this approach precludes our understanding of fluctuating interactions and competition between defense systems, as well as the consequences and trade-offs of phage resistance. Furthermore, the extent to which a given defense system is expressed in its native context as well as the potential for dissemination of resistance to naïve hosts through horizontal transfer cannot be assessed. These approaches also do not support the discovery of phage-encoded counter defenses, which lag behind our knowledge of anti-phage defenses but are likely key drivers of defense diversification.

A promising model system to address unanswered questions in phage-host ecology and evolution lies in the long-term sampling of *Vibrio cholerae* and its lytic phages (vibriophages). Pathogenic *V. cholerae* strains of the O1 serogroup possessing the key virulence factor cholera toxin are responsible for the infectious diarrheal disease cholera, which continues to pose a significant global public health threat (15). In Bangladesh, where cholera is endemic, lytic phage predation is implicated in modifying the duration and severity of cholera outbreaks (16, 17). Interestingly, cholera patients can shed both vibriophages and viable *V. cholerae*, indicating that complex interactions between phages and their hosts take place within the human intestine, as neither entity is driven to extinction. Long-term sampling of cholera-infected patients in Bangladesh has revealed the persistence of three dominant lytic phages ICP1, ICP2 and ICP3 (18), making this a powerful system to understand biologically relevant outcomes of phage predation and to discover reciprocal adaptations in phages infecting pathogenic bacteria within the context of human disease.

### SXT ICEs determine clinical *Vibrio cholerae* phage susceptibility

To gain insight into the ongoing co-evolutionary dynamics between *V. cholerae* and its phages, we evaluated whether clinical *V. cholerae* isolates were susceptible to infection by ICP1 from past, future or contemporaneous patient samples (Fig. 1A). A pattern emerged in which *V. cholerae* isolates were uniformly susceptible to contemporaneous phage, but restricted ICP1 from temporally mismatched patients (Fig. 1B). *V. cholerae* isolates from the past were susceptible to phages from this time period, but restricted infection of phage from future patient samples, while isolates from 2018 onward restricted phages from earlier sampling dates but were susceptible to contemporaneous phages (Fig. 1B). Importantly, *V. cholerae* DCP66 from December 2017 was uniquely susceptible to all ICP1 isolates, suggesting it lacks a phage resistance determinant, and phages from this sample were uniquely restricted by all other isolates of *V. cholerae*.

**Fig. 1.**
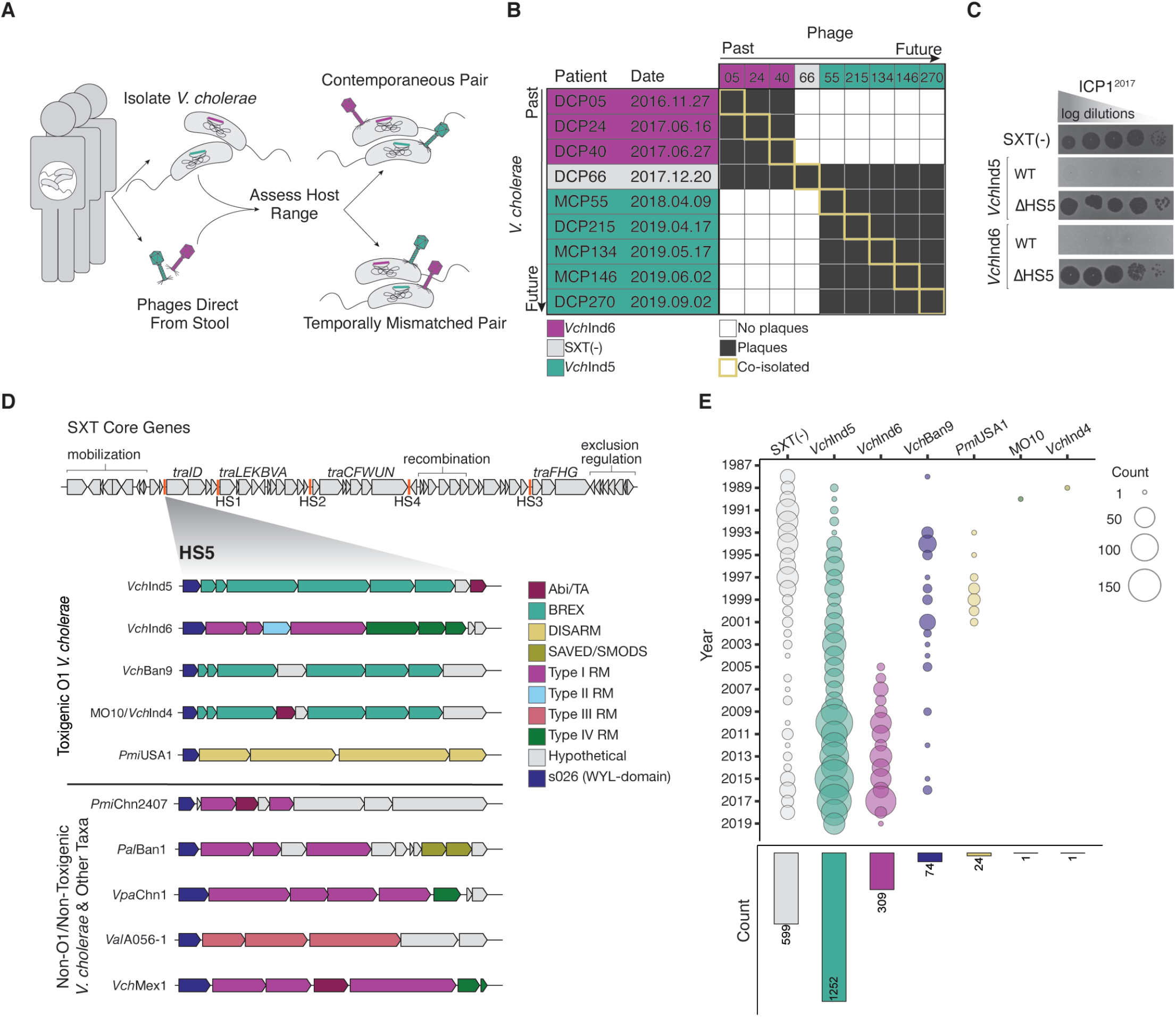
The prevalence and identity of SXT ICEs in clinical *V. cholerae* determines susceptibility to phages. (A) Schematic of time-shift experiments. ICP1 phage in cholera patient stool samples were probed for ability to productively infect isolates of *V. cholerae* from contemporaneous samples or from the past or future. Matching colors denote contemporaneously isolated *V. cholerae* and phage pairs. (B) *V. cholerae* (vertical axis) susceptibility matrix to ICP1 phages from past or future patients (top horizontal axis), where colors denote the presence of the SXT ICE (as identified through whole genome sequencing of patient isolates) in *V. cholerae* from the patient stool sample. ICE*Vch*Ind6 *V. cholerae* were susceptible to phages shed from patients in which ICE*Vch*Ind6 *V. cholerae* was recovered, but restricted phages from patients in which SXT(−) or ICE*Vch*Ind5 *V. cholerae* was recovered. The SXT(−) isolate was only susceptible to co-isolated phages. ICE*Vch*Ind5 *V. cholerae* isolates restricted phages from the past but were susceptible to phages from other patients in which ICE*Vch*Ind5 *V. cholerae* was recovered. (C) Tenfold dilutions of ICP1^2017^ (isolated from patient DCP24, propagated on SXT(−) *V. cholerae*) spotted on an SXT(−) *V. cholerae* host or exconjugant with the SXT ICE indicated (bacterial lawns in gray, zones of killing are shown in black). (D) Genomic organization of SXT ICE core genes (core function labeled above, *tra* genes encode functions related to conjugation) and hotspot regions (HS, denoted by orange lines) (modified from Wozniak et a. 2009) (not to scale). Diverse anti-phage systems are encoded in hotspot 5 of SXT ICEs found in toxigenic O1 *V. cholerae* (top) and other hosts (bottom). All hotspot 5 regions shown have a 3’ WYL-domain protein (Fig. S1), previously shown to be associated with defense systems (24). (E) Temporal distribution of SXT ICEs in toxigenic O1 *V. cholerae* between 1987-2019 (n=2,260). The size of the circle scales with the number of genomes analyzed per year, with total counts for each SXT ICE denoted below.

Differences in *V. cholerae’s* susceptibility to ICP1 could not be explained by known anti-phage systems. However, comparative genomics of clinical *V. cholerae* isolates from this time period revealed the variable presence and identity of an integrative and conjugative element (ICE) of the SXT/R391 family (19). This ~100kb island is of clinical importance for pathogens such as *Proteus mirabilis* and *V. cholerae,* conferring antibiotic resistance to <u>s</u>ulfametho<u>x</u>azole, <u>t</u>rimethoprim (SXT) and, variably, streptomycin, tetracycline, kanamycin and chloramphenicol (20, 21). SXT ICEs are composed of core and accessory genes, with the core genes encoding functions related to SXT regulation and transmission through conjugative transfer (Fig. 1D). Accessory genes, such as those conferring resistance to heavy metals or antibiotics, determine the identity of the SXT ICE and are clustered in five “hotspot” regions as well additional variable regions in subset of SXT ICEs (20) (Fig. 1D). The temporal pattern of phage resistance observed during this sampling period correlated with the transition of ICE*Vch*Ind6(+) to SXT(−) to ICE*Vch*Ind5(+) *V. cholerae*, suggesting that SXT ICEs may confer protection against clinically relevant vibriophages. To test whether SXT ICEs can confer phage resistance, we conjugated ICE*Vch*Ind6 and ICE*Vch*Ind5 into the phage sensitive *V. cholerae* SXT(−) strain E7946. Each SXT ICE was sufficient to inhibit plaque formation by ICP1 recovered during the sampling period (Fig. 1C). These results demonstrate that SXT ICEs found in *V. cholerae* natively express a resistant determinant sufficient to restrict vibriophage co-circulating with SXT ICEs in cholera patients.

The genes predicted to be responsible for phage resistance in ICE*Vch*Ind5 and ICE*Vch*Ind6 were readily identified, and in both ICEs the genes localized to hotspot 5. ICE*Vch*Ind5 hotspot 5 contains a type 1 bacteriophage exclusion (BREX) system, an abundant phage resistance system that restricts phages through an unknown mechanism known to rely on epigenetic modification (14, 22). Additionally, ICE*Vch*Ind5 hotspot 5 encodes a putative abortive infection gene (Abi), and a gene of unknown function (Fig. 1D). ICE*Vch*Ind6 contains several restriction-modification (RM) systems as well as hypothetical genes in hotspot 5 (Fig. 1D). Deletion of hotspot 5 from ICE*Vch*Ind5 and ICE*Vch*Ind6 rendered *V. cholerae* susceptible to ICP1 infection, demonstrating that the anti-phage systems in hotspot 5 are necessary for phage resistance (Fig. 1C). On the basis of these results, we examined hotspot 5 regions in all published SXT ICEs by supplementing data from ICEBerg 2.0 (23) with additional published SXT ICEs since the database was last updated. Strikingly, we observed that all 76 de-duplicated SXT ICE genomes published to date harbor predicted anti-phage systems in the same hotspot 5 location, with the exception of three elements that had no discernible hotspot 5, suggesting it was lost through recombination. All hotspot 5 regions share a putative transcriptional regulator possessing a WYL-domain previously shown to associate with defense systems (24) (Fig. 1D). A phylogenetic tree based off of the nucleotide sequence of the WYL-domain gene clustered based on the type of anti-phage system encoded in hotspot 5, and builds a tree that does not agree with the relationship inferred from the conserved gene downstream of hotspot 5 *traI* (Fig. S1). This demonstrates that gene cargo dedicated to phage defense is a ubiquitous yet unrecognized facet of the biology of SXT ICEs (Fig, 1D). Hence, although SXT ICEs are infamous for their role in the spread of antibiotic resistance, these elements also encode for and thus disseminate phage resistance throughout Gammaproteobacteria, including in human pathogens like *Proteus mirabilis* and *V. cholerae* for which phage therapy/prophylaxis has been proposed (25–27).

To ascertain the global relevance of SXT ICE-mediated phage resistance in *V. cholerae*, we determined the identity and prevalence of SXT ICEs in published toxigenic O1 *V. cholerae* genomes, and in a collection of 252 isolates sequenced as a part of this study. Genomes of 2,260 isolates were analyzed with isolation dates going back to 1987 when SXT was first identified in toxigenic *V. cholerae* (28), to late 2019 (Fig. 1E). Previous analysis of 96 SXT ICEs from seventh pandemic *V. cholerae* isolates identified a total of five different SXT ICEs in toxigenic *V. cholerae*, and found that ICE*Vch*Ind5 dominated globally (29). Our expanded analysis demonstrates that ICE*Vch*Ind5 remains globally dominant, followed by ICE*Vch*Ind6 and ICE*Vch*Ban9 (also known as ICE*Vch*Moz10) (Fig. 1E). Additional SXT ICEs are found sporadically in toxigenic *V. cholerae*, including MO10 (the first discovered and best characterized SXT ICE) and ICE*Vch*Ind4 (also known as ICE*Vch*Ban11) (Fig. 1E). Of note, we found an additional SXT ICE, ICE*Pmi*USA1, not previously described in *V. cholerae* which contains genes in hotspot 5 with PFAM domains characteristic of the DISARM system (13) (Fig 1D and 1E). Despite the obvious selective advantage afforded by SXT ICEs in conferring antibiotic resistance in *V. cholerae*, we also find SXT(−) toxigenic O1 strains persist through this sampling period.

### Genetic dissection of SXT ICE hotspot 5-mediated restriction of phages and MGEs

To gain further insight into how SXT ICEs influence *V. cholerae* ecology and gene flux, we tested the ability of SXT ICEs to restrict foreign genetic elements that are clinically important to *V. cholerae*. We focused on the three most prevalent SXT ICEs in *V. cholerae* for further study, ICE*Vch*Ind5, ICE*Vch*Ind6 and ICE*Vch*Ban9, which contain either a BREX system, or several restriction-modification systems in hotspot 5 (Fig. 2A). *V. cholerae* exconjugants possessing each SXT ICE were challenged with the three dominant lytic vibriophages found in cholera patient stool samples: ICP2, ICP3 (both *Podoviridae*) and two temporally distinct isolates of ICP1 (*Myoviridae*). Each SXT ICE tested conferred resistance to some, but not all, of the vibriophages tested (Fig. 2B, Fig. S2). Although ICE*Vch*Ind5 and ICE*Vch*Ban9 both encode type 1 BREX systems, they showed different restriction profiles with regards to ICP1, likely because they possess highly divergent genes encoding for BrxX (26% amino acid identity), the essential methyltransferase and restriction determinant for BREX(14).

**Fig. 2.**
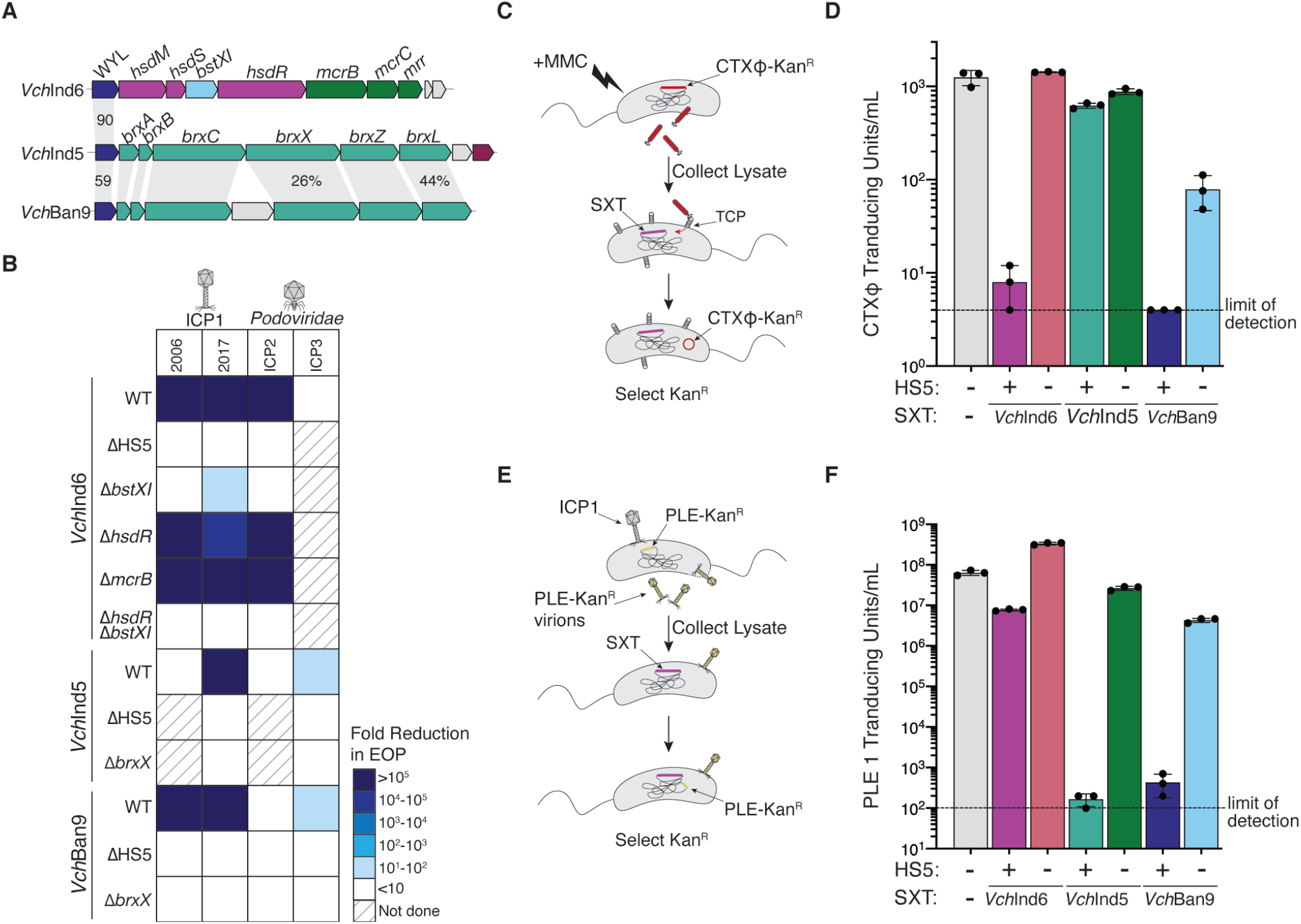
SXT ICEs confer resistance to phages and mobile elements in *V. cholerae*. (A) Genomic organization of hotspot 5 of the three dominant SXT ICEs in toxigenic O1 *V. cholerae* labeled according to homologous genes identified by RE Base/PFAM searches. Color scheme is identical to Fig. 1D. Shading denotes proteins with same predicted function; Percent identity is indicated for genes encoding proteins that share >20% amino acid identity, if unlabeled percent identity is <20%. (B) Fold reduction in the efficiency of plaquing (EOP) of lytic vibriophages afforded by SXT ICEs under native expression levels. When the SXT ICE did not reduce EOP, deletions were not tested. Source data is shown in Fig. S2. (C) Schematic representation of the filamentous prophage CTXϕ transduction assay. (D) Transduction of antibiotic marked CTXϕ into *V. cholerae* possessing each SXT ICE and derivatives with hotspot 5 deletions. (E) Schematic representation of PLE transduction. (F) Transduction of antibiotic marked PLE 1 into *V. cholerae* possessing each SXT ICE and derivatives lacking hotpot 5.

We next wanted to determine which system is responsible for phage inhibition in *V. cholerae*, and whether different restriction enzymes restrict different phages. From ICE*Vch*Ind6, each restriction enzyme was deleted, while the methyltransferase *brxX* was deleted from ICE*Vch*Ind5 and ICE*Vch*Ban9. ICE*Vch*Ind5Δ*brxX* and ICE*Vch*Ban9Δ*brxX* no longer inhibited ICP1 and ICP3, showing that the BREX system is responsible for inhibition of lytic vibriophages (Fig. 2B). Deletion of each putative restriction enzyme in ICE*Vch*Ind6 showed that the protein with homology to BstXI (protein ID <u>AIM16524.1</u>), a type II restriction enzyme, is largely responsible for ICE*Vch*Ind6’s anti-phage activity, however, there were differences between ICP1 isolates (Fig. 2B). Specifically, ICE*Vch*Ind6Δ*bstXI* was susceptible to ICP1^2006^, yet ICP1^2017^ was still partially inhibited. Deletion of the type I restriction enzyme *hsdR* also had differential effects on the two isolates of ICP1 tested. ICE*Vch*Ind6Δ*hsdR* still completely inhibited ICP1^2006^, but ICP1^2017^ was able to spontaneously overcome ICE*Vch*Ind6Δ*hsdR* with plaquing that did not scale with the dilution series (Fig. S3), suggesting that HsdR has some activity against ICP1^2017^, which notably co-circulated with ICE*Vch*Ind6(+) *V. cholerae*.

Next, we asked whether SXT ICEs restrict other MGEs, including temperate phage that can provide beneficial traits to their hosts upon integration into the bacterial chromosome. Specifically, we assessed whether SXT ICEs restrict acquisition of two clinically important and beneficial MGEs in epidemic *V. cholerae*: the prophage CTXϕ and the phage-inducible chromosomal island like elements (PLEs). The filamentous phage CTXϕ encodes cholera toxin, and is essential for *V. cholerae’s* pathogenicity in the human host (30). PLEs are ~20kb phage satellites that confer resistance to ICP1 and have been a persistent part of the mobilome of epidemic *V. cholerae* dating back to at least 1949, before SXT was common in clinical *V. cholerae* isolates (31). The acquisition of antibiotic marked CTXϕ and each PLE was assessed in otherwise isogenic *V. cholerae* hosts either lacking an SXT ICE or harboring each of the most prevalent SXT ICEs (ICE*Vch*Ind6, ICE*Vch*Ind5, ICE*Vch*Ban9), or derivatives lacking hotspot 5 (Fig. 2C and 2E). SXT ICEs restricted acquisition of both PLEs and CTXϕ, but to varying extents, and in all cases, transduction was restored in mutants lacking hotspot 5 (Fig. 2D and 2F). PLE 1 was uniquely restricted by ICE*Vch*Ind6 compared to PLEs 2, 3, 4 and 5 (Fig 2F and Fig. S4B-E). The difference in inhibition of different PLEs can be explained by the predicted recognition motif for BstXI from *Bacillus subtilis*, which shares 50% amino acid identity with ICE*Vch*Ind6’s BstXI homolog: CCANNNNN^NTGG ((32)), which is found once in PLE 1 but not in other PLEs. Knocking out this site restored PLE 1 transduction to the level observed for the SXT(−) host (Fig. S4A), again demonstrating the relative contribution of BstXI to ICE*Vch*Ind6’s restriction of foreign elements. Collectively these data demonstrate that like other anti-phage defense systems, SXT-mediated restriction of MGEs comes at a cost. While all SXT ICEs restricted infection by clinically relevant lytic vibriophages, they also limited acquisition of CTXϕ and PLEs, which provide beneficial traits to epidemic *V. cholerae*, observations that help to explain the maintenance of SXT(−) strains in nature (Fig. 1E).

### SXT ICEs disseminate phage resistance to other Gammaproteobacteria

SXT ICEs are recombinogenic, self-transmissible mobile elements hypothesized to evolve by genetic exchange of variable and hotspot regions (29, 33) suggesting that gene cargo in hotspot 5 could be mobilized between taxa through both SXT ICE conjugation and through recombination of hotspot 5 with other SXT ICEs or homologous DNA. To determine the extent of lateral dissemination, genes in hotspot 5 from ICE*Vch*Ind6, ICE*Vch*Ind5 and ICE*Vch*Ban9 were queried using BLASTn to probe whether the genes could be found in other taxa, either within SXT ICEs or in a different genetic context. The genes in ICE*Vch*Ind6 hotspot 5 were always localized to an SXT ICE, and ICE*Vch*Ind6 in its entirety is found in other Gammaproteobacteria with over 99% nucleotide identity (Fig. 3A, Table S3). Intriguingly, a variant of ICE*Vch*Ind6 was also identified in *Pseudoalteromonas* sp. referred to as ICE*Psp*Spa1(34). ICE*Psp*Spa1 is 96.45% identical to ICE*Vch*Ind6 except in place of *bstXI* is a SIR2 domain protein that is implicated in the Thoeris defense system (11) (Fig. 3A). Despite being globally dominant in toxigenic *V. cholerae* O1 strains, the ICE*Vch*Ind5 BREX system was rarely found outside of *V. cholerae*, occurring in hotspot 5 from an otherwise divergent SXT ICE in *P. mirabilis* and outside of an MGE in *Shewanella* (Fig. 3A). The BREX system from ICE*Vch*Ban9 was the most widely distributed, both taxonomically and in terms of genetic context, being found within SXT ICEs (for example, in *Escherichia coli* and *P. mirabilis*), as well as outside of SXT ICEs. These varied locations include the chromosome, on plasmids, as well as in non-SXT ICEs (Fig. 3A and Table S3-S5). The distribution of hotspot 5 encoded anti-phage systems in other Gammaproteobacteria suggests that these systems confer resistance to a broad range of phages infecting different hosts well beyond *V. cholerae,* and that the self-transmissible nature of SXT ICEs would disseminate these anti-phage systems to new taxa.

**Fig. 3.**
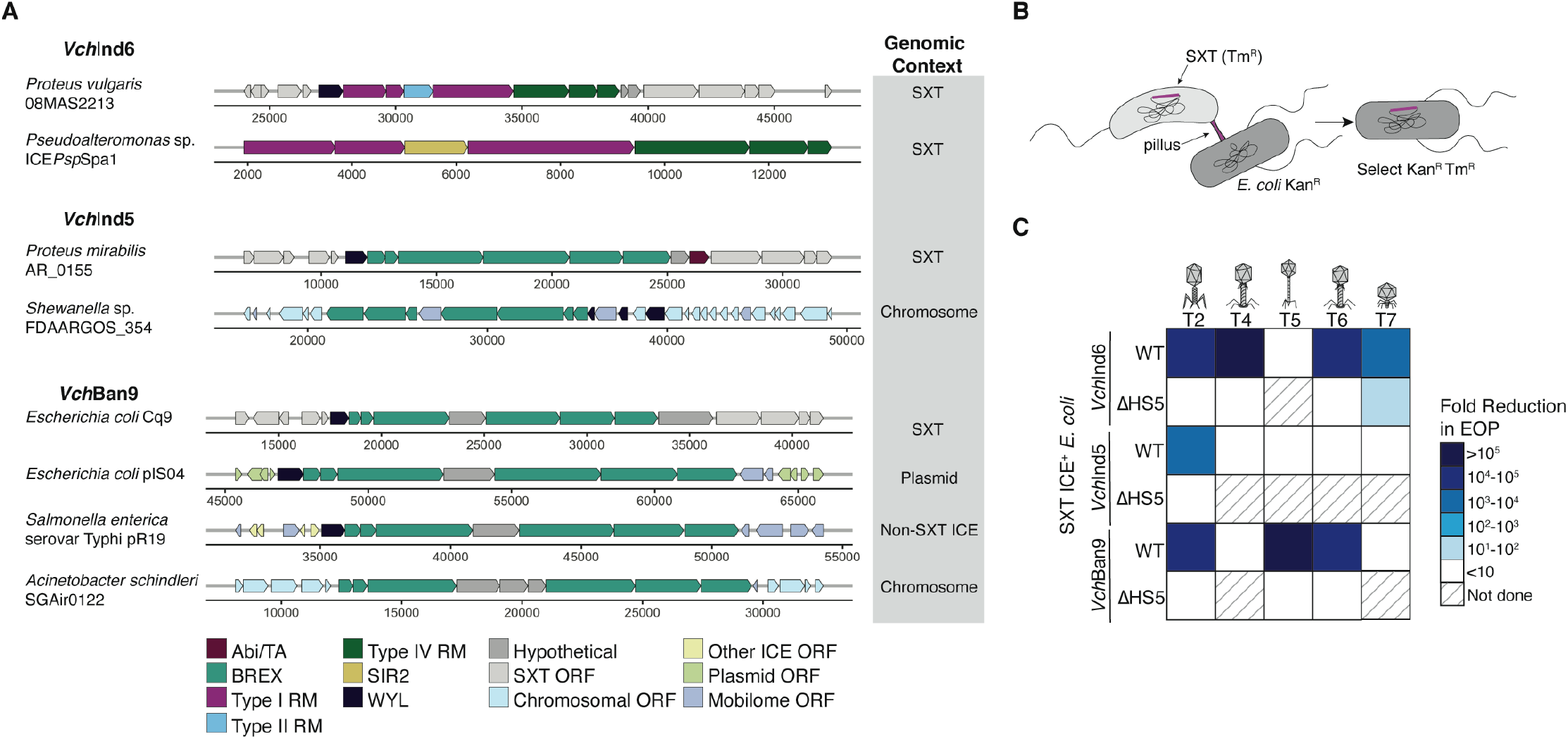
Hotspot 5 from *V. cholerae* SXT ICEs occur in other Gammaproteobacteria and confer phage resistance upon conjugation into *E.coli*. (A) Gene maps depicting instances of hotspot 5 genes (from the start of the WYL-domain protein to the end of the last gene) from ICE*Vch*Ind6, ICE*Vch*Ind5 and ICE*Vch*Ban9 found outside of *V. cholerae*, identified using BLASTn (shown are genes with >90% nucleotide identity across >75% of the query). The genomic context of the hotspot 5 genes identified is indicated in the shaded box to the right. (B) Schematic depicting SXT ICE conjugative transfer from *V. cholerae* into *Escherichia coli* K12 MG1655. (C) Fold-reduction in the efficiency of plaquing (EOP) of lytic coliphages mediated by SXT ICEs and corresponding hotspot 5 deletions in *E. coli*, under native expression levels. When the SXT ICE did not reduce EOP, the deletion was not tested. Source data is shown in Fig. S5.

The test whether SXT ICEs can confer phage resistance upon entry into a new host, ICE*Vch*Ind6, ICE*Vch*Ind5 and ICE*Vch*Ban9 and the cognate Δhotspot 5 derivatives, were conjugated into *E. coli* (Fig. 3B) and exconjugants were challenged with the lytic coliphages T2, T4, T5, T6 and T7. Strikingly, all SXT ICEs tested offered protection from at least one coliphage, with the broadest protection afforded by ICE*Vch*Ind6 (restricting all phage except for T5) and the narrowest restriction afforded by ICE*Vch*Ind5 (which only partially restricted T2) (Fig. 3C, Fig. S5). ICE*Vch*Ban9, which is natively found in *E. coli* (Fig 3A), restricted T2, T5 and T6, while T4 and T7 were not inhibited. The T7-encoded DNA mimic protein, Ocr, was recently shown to inhibit type 1 BREX (35) and thus may explain T7’s resistance to ICE*Vch*Ban9 and ICE*Vch*Ind5. *E.coli* exconjugants harboring the Δhotspot 5 SXT ICE derivatives were susceptible to all phages tested, demonstrating that it is the same anti-phage systems in hotspot 5 that confers resistance in *E.coli* and *V. cholerae*. This demonstrates that upon acquisition of SXT ICEs through conjugation, hotspot 5 is natively expressed in the recipient taxa and confers resistance to their cognate phage.

### Co-Evolution of Vibriophage ICP1 with *Vch*Ind5

In light of the discovery that SXT(+) *V. cholerae* restricts phages through anti-phage systems known to rely on differential epigenetic modification and subsequent restriction, we postulate that phages shed from patients infected with SXT ICE(+) *V. cholerae* have escaped restriction through stochastic acquisition of DNA modification. Epigenetically modified phage can then productively infect co-circulating strains possessing the same SXT ICE, but are unable to infect *V. cholerae* with a different SXT ICE (as shown in our time-shift assay, Fig. 1B). In further support of this, phage in the patient stool in which we recovered SXT ICE(−) *V. cholerae* DCP66 were restricted by all SXT ICE(+) hosts, and the SXT ICE(−) *V. cholerae* isolate from this patient was uniquely susceptible to infection by phages from all patient stool samples (Fig. 1B). Other genetic means of overcoming modification-based restriction are common among phages and include anti-restriction proteins, modification of DNA bases and loss of recognition motifs (reviewed in (36). To distinguish between potential mechanisms of epigenetic or genetic escape, ICP1 isolates from patient stool samples were first passaged on SXT ICE(−) *V. cholerae*, and then evaluated for the ability to productively infect the clinical *V. cholerae* strains from our original time-shift experiment (Fig. 4A). ICP1 isolates from stool samples in which we co-recovered ICE*Vch*Ind6(+) *V. cholerae* lost the ability to productively infect isolates with ICE*Vch*Ind6, suggesting that ICP1 shed in patient stool samples had escaped through acquiring modification during the course of infection (Fig. 4A). In contrast, the pattern of susceptibility seen with ICE*VchI*nd5 and cognate phages indicates a genetic route of escape. Specifically, ICP1 shed from patient stool samples with *V. cholerae* harboring ICE*Vch*Ind5 retained the ability to infect co-circulating ICE*Vch*Ind5(+) isolates after passage through an SXT(−) host, yet ICP1 isolates shed in patient samples in which we recovered ICE*Vch*Ind6(+) *V. cholerae* were still restricted by ICE*Vch*Ind5 (Fig. 4A). This was reminiscent of the phenotype for ICP1^2006^, which was not restricted by ICE*VchI*nd5 (Fig. 2B). To identify the genetic mechanism responsible for the host range differences between ICP1 isolates, we compared the genomes of ICP1 isolates from across a temporal gradient that can and cannot infect *V. cholerae* harboring ICE*Vch*Ind5 (Fig. 4B). This analysis revealed differences in the operon encoding gene products (gp) 21-25 that co-varied with plaque formation on a ICE*Vch*Ind5(+) host (Fig. 4B and 4C). Unlike ICP1 isolates that can productively infect ICE*Vch*Ind5(+) *V. cholerae*, the restricted ICP1^2017^ lacks both *gp21* and the promoter region for *gp25*, which we identified by analyzing the ICP1 transcriptome (Barth et al., 2020) (Fig. 4C), suggesting one of these early genes may allow ICP1 to overcome restriction by ICE*Vch*Ind5. Having narrowed in on these two gene products as candidate anti-ICE*Vch*Ind5 proteins, we evaluated if *in trans* expression of either *gp21* or *gp25* was sufficient to rescue plaque formation for the restricted ICP1^2017^ phage. Ectopic expression of *gp25*, but not *gp21*, restored ICP1^2017^’s ability to form plaques on *V. cholerae* harboring ICE*Vch*Ind5 (Fig. 4D), supporting our designation of *gp25* as *orbA* for “overcome restriction by BREX”. To decipher if OrbA was the sole factor permitting ICP1 plaque formation on the ICE*Vch*Ind5(+) host, we expressed *orbA* during ICP3 infection, since ICP3 is an unrelated phage that is also restricted by the BREX system in ICE*Vch*Ind5 (Fig. 2B). Here too we observed that *orbA* expression restored productive infection for ICP3 on the ICE*Vch*Ind5(+) host (Fig. 4D). These results demonstrate that OrbA (YP_004250966) functions as an anti-BREX protein with activity against ICE*Vch*Ind5. OrbA encodes a small 44 amino acid protein with no homology to any annotated proteins, as well as no PFAM/DUF domains. Furthermore, PSI-BLAST did not return any significant homologs outside of ICP1. The T7-encoded anti-restriction protein Ocr has been shown to overcome BREX in *E. coli* by functioning as DNA mimic (35). Although the crystal structure for OrbA has not been solved, a predicted model does not show the electrostatics characteristic of a DNA mimic like Ocr (Fig. S6), suggesting a different mechanism of action for OrbA’s anti-BREX activity. Our findings also indicate that OrbA is not a broadly active anti-BREX protein since the BREX system encoded by ICE*Vch*Ban9 equally restricts ICP1^2006^ and ICP1^2017^ (Fig. 2B).

**Fig. 4.**
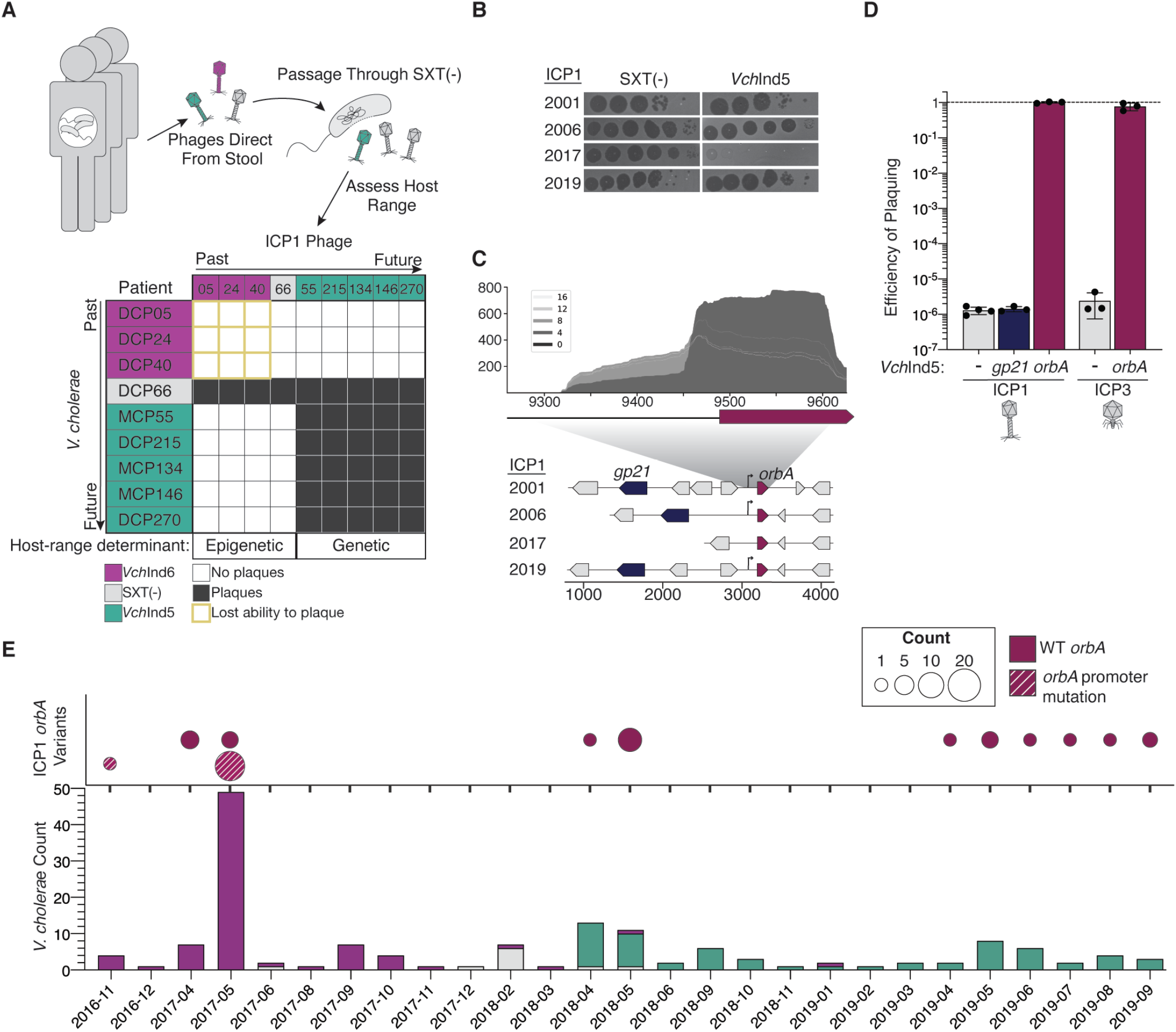
Clinical ICP1 overcomes co-circulating SXT ICEs through acquisition of epigenetic modification as well as an anti-BREX protein ObrA. (A) ICP1 isolates from cholera-patient stool samples were propagated on an SXT(−) strain and then probed for ability to infect clinical *V. cholerae* isolates. ICP1 isolates from ICE*Vch*Ind6(+) stool samples lost the ability to infect co-circulating *V. cholerae* upon passage through an SXT(−) host indicative of initial escape through epigenetic modification. ICP1 isolates from ICE*Vch*Ind5(+) patients retained infectivity of co-circulating *V. cholerae* after passage through an SXT(−) host, suggestive of a genetic means of overcoming ICE*Vch*Ind5. (B) Tenfold dilutions of ICP1 isolates spotted on SXT(−) *V. cholerae* or an isogenic exconjugant of ICE*Vch*Ind5 identified ICP1^2017^ (from patient DCP24) as uniquely inhibited by ICE*Vch*Ind5 (bacterial lawns in gray, zones of killing are shown in black). (C) Comparative genomics of ICP1 isolates showing that ICP1^2017^ lacks *gp21*-*24* and the promoter for *gp25*, as identified through RNA-seq of ICP1^2006^ (top, displaying average read coverage over the course of infection, reads are color coded by time point in minutes post infection). (D) Ectopic expression of *gp21* and *gp25* in ICE*Vch*Ind5(+) *V. cholerae*: *gp25* (designated *orbA* for “overcome restriction by BREX”) but not *gp21* restores plaquing by ICP1^2017^ as well the unrelated lytic phage ICP3. (E) Sampling and whole genome sequencing of 152 *V. cholerae* isolates and 44 ICP1 isolates from cholera patient stool samples in Bangladesh, depicting isolate counts (y-axis) over time, highlighting the transition from ICE*Vch*Ind6 to SXT(−) to ICE*Vch*Ind5(+) *V. cholerae*. ICP1 isolates with the *gp25* promoter mutation were only isolated from patients in which we recovered ICE*Vch*Ind6(+) *V. cholerae*, and have not been isolated since the re-emergence of ICE*Vch*Ind5(+) *V. cholerae*.

We next set out to document the reciprocal dynamics between the repertoire of SXT ICEs in *V. cholerae* with anti-BREX activity in ICP1 (afforded by OrbA) in greater detail. Towards this end, we performed whole genome sequencing of 152 toxigenic O1 *V. cholerae* isolates and 44 ICP1 phages isolated from cholera patient stool samples in Bangladesh between November 2016 and September 2019. As was evident in the subset of samples in which ICP1 and *V. cholerae* were co-isolated (Fig. 1B), we observed a transition between circulating ICE*Vch*Ind6(+) strains to ICE*Vch*Ind5(+) strains in 2018, with intermittent recovery of SXT ICE(−) *V. cholerae* from patients between June 2017 and May 2018. Interestingly, ICP1 isolates lacking the *orbA* promoter region (as in ICP1^2017^ (Fig. 4C)) were only isolated during a time period when ICE*Vch*Ind5 was absent in cholera patients (Fig. 4E); however, concurrent with the emergence of *V. cholerae* ICE*Vch*Ind5(+) in patient stool samples in the spring of 2018, all contemporary ICP1 isolates possess the promoter for *orbA*. This locus provides an example of selection for a novel anti-BREX gene found in phage, which can counter restriction by an SXT ICE, and shows how temporal dynamics in SXT ICEs can exert purifying selection on phages. In times when ICP1 is not encountering ICE*Vch*Ind5 in nature, the locus that confers resistance to ICE*Vch*Ind5 is not under positive selection. Thus, variants with promoter mutations persist, or are perhaps selected for, suggesting this locus provides no fitness benefit to ICP1 when it is not co-circulating with ICE*Vch*Ind5(+) *V. cholerae*.

### Phage infection stimulates SXT ICE conjugation

SXT ICEs are SOS responsive, meaning that DNA damage leading to the accumulation of single stranded DNA triggers de-repression of SXT, culminating in SXT excision and expression of operons related to conjugative transfer (38). DNA damaging agents such as mitomycin C (MMC), ultraviolet radiation, and certain classes of antibiotics that interfere with DNA replication (for example, ciprofloxacin), have been shown to increase SXT conjugation frequency (39). DNA damage may serve as a proxy for impaired cell viability, thus SXT ICEs that can mobilize and spread horizontally to escape a compromised host would have a selective advantage. We postulated that cell damage from infection by lytic phages could serve as an input into SXT ICE de-repression and transfer. To test this hypothesis, we evaluated the frequency of conjugative transfer of ICE*Vch*Ind5 into recipient *V. cholerae* following infection by either ICP1^2006^ or ICP1^2017^, representing phages that are not restricted or are fully restricted by ICE*Vch*Ind5, respectively (Fig 2B). Overnight cultures of ICE*Vch*Ind5(+) *V. cholerae* were infected with ICP1 at a multiplicity of infection of 0.5, and frequency of transfer into a differentially marked *V. cholerae* recipient was determined. Importantly, the added differentially marked *V. cholerae* recipient lacks the O1 antigen, which is the receptor for ICP1 (18), making the recipient insensitive to ICP1 infection, yet proficient for conjugation (Fig. S7C). As expected, conjugation frequency was elevated above background following treatment with only MMC (Fig. 5A). Interestingly, infection by ICP1^2006^ (but not the restricted phage ICP1^2017^) resulted in high frequency SXT ICE transfer compared to an uninfected control, comparable to conjugation frequency observed from a MMC treated donor (Fig. 5A). This increase in conjugation frequency per donor cell was driven by a consistent frequency of transfer despite productive phage infection among the donor strains resulting in a 100-fold decrease in donor cell viability, yet remarkably, constant transfer frequency (Fig. S8). As expected, no antibiotic resistant exconjugants were detected with a Δ*traD* donor, demonstrating that phage-mediated transduction is not a contributing factor in these assays (Fig. 5A). It is unclear whether it is the infected cells that are conjugating at a greater frequency, or if the uninfected cells in the population are sensing phage lysis through an unknown mechanism and becoming high frequency donors. We propose alternative models, the first where cellular damage wrought by phage infection triggers increased transfer of SXT ICEs as the SXT ICE races to transfer out of a dying host (Fig. 5B), or, alternatively, productive phage infection in neighboring cells leads to increased conjugative transfer from uninfected cells, through a yet undescribed mechanism.

**Fig. 5.**
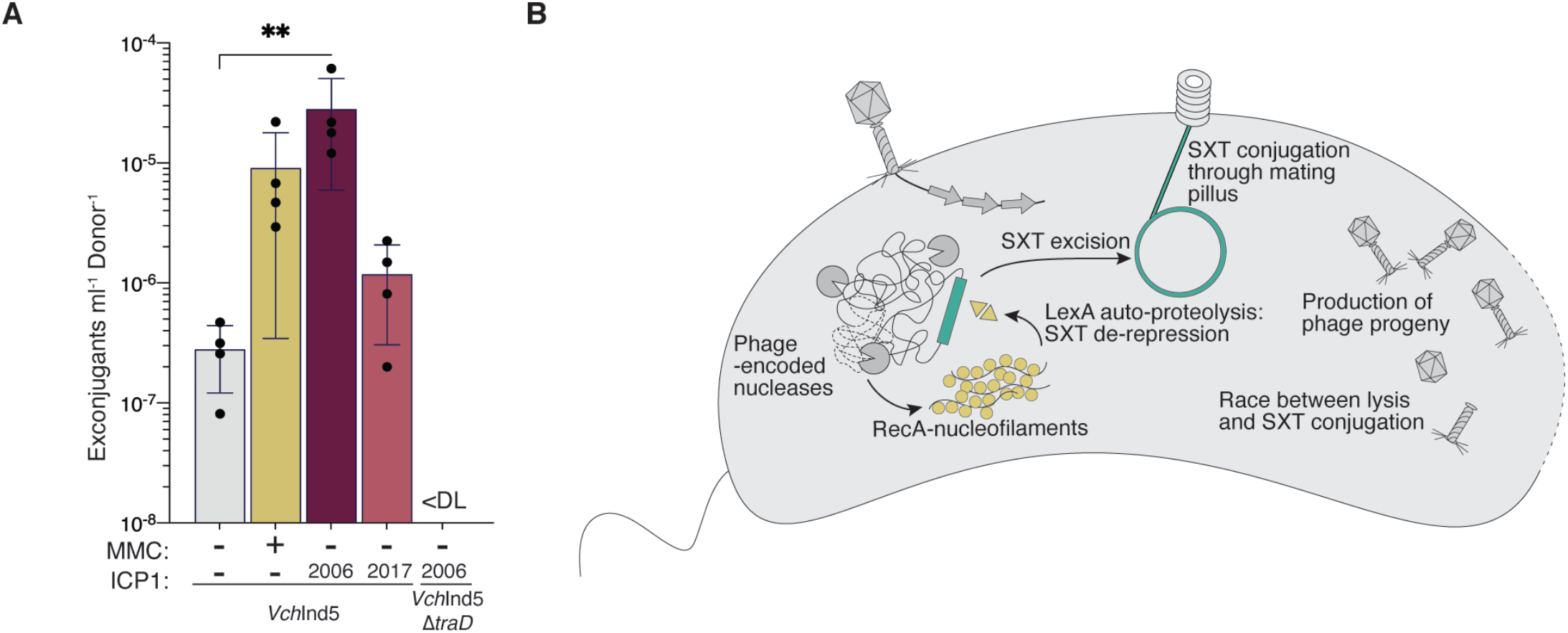
ICP1 infection increases SXT ICE conjugation frequency. (A) Conjugation frequency from a donor *V. cholerae* mock treated (LB), mitomycin C (MMC) treated or infected by ICP1^2006^ (unrestricted phage) or ICP1^2017^ (restricted phage). Asterisks denotes significance by ANOVA p-value <0.01. No exconjugants were detected from a phage infected donor lacking *traD* (dashed line indicates limit of detection). (B) Model representing a possible mechanism of ICP1-stimulated SXT conjugation: Upon infection, ICP1-encoded nucleases degrade the *V. cholerae* genome, triggering SXT de-repression through the SOS response, resulting in the SXT ICE escaping the lysing host cell through conjugation.

## Discussion

One of the prominent genomic features of current seventh pandemic *V. cholerae* are ICEs of the SXT/R391 family, infamous for conferring multidrug resistance (19, 20, 29). SXT ICEs appeared in sequenced epidemic *V. cholerae* in 1987 and currently dominate in clinical isolates in regions where cholera is endemic (28, 40, 41). Here we show that SXT ICEs invariably encode phage-resistance genes in hotspot 5, linking together antibiotic and phage resistances on a single mobile genetic element widespread in Gammaproteobacteria. This builds on a previous study which showed the SXT ICE*Vsp*Por3 reduced plaquing by the phage T1 when conjugated into *E. coli* (42), however, it was not demonstrated that this was due to hotspot 5, or whether ICE*Vsp*Por3 offered protection against cognate phages in its native host context.

Co-evolution between *V. cholerae* and lytic vibriophages is driven, at least in part, by presence and identity of SXT ICEs. Using a time-shift experiment (43), we find that lytic phage can infect contemporary *V. cholerae* hosts but cannot infect hosts from the past or the future. This pattern is consistent with negative frequency-dependent dynamics, where the faster evolving partner (here, lytic phage) has higher fitness in the presence of the contemporary antagonist than those from the past or the future (44–46). However, over a short time scale, co-evolution of ICP1 with *V. cholerae* appears to be driven by selective sweeps. Adaptations that allow for phage infection of contemporary SXT ICE(+) *V. cholerae* sweep to fixation over short times scales, rendering contemporary ICP1 more fit that ICP1 from the near past, which is characteristic of arms race dynamics (45–47). ICP1’s adaptations to overcome contemporary SXT ICEs is driven both by changes in its capacity to express the anti-ICE*Vch*Ind5 BREX gene product *orbA* and acquisition of epigenetic modification allowing for infection of ICE*Vch*Ind6 hosts. The heterogeneity in ICE SXT identity and presence in circulating toxigenic *V. cholerae* suggests the potential for population bottlenecks and subsequent sweeps. Phages that have acquired anti-SXT ICE mechanisms will sweep through the population, providing a selective advantage to hosts with SXT ICEs that are resistant to the dominant phages. The subsequent sweep of the resistant SXT ICE through the *V. cholerae* population generates a further bottleneck on the ICP1 population. These dynamics of fluctuating genomic dominance in a population of bacteria and phages allows for apparent long-term coexistence between phages and their bacterial hosts, where neither entity drives the other to extinction. The heterogeneity in SXT ICE-mediated phage defense can be considered a “pan-immune system”, where the collective pool of SXT ICEs circulating in *V. cholerae* serve to restrict all lytic phages: ICP1, ICP2 and ICP3 (2). Additionally, these temporal dynamics provide insight into the ongoing maintenance of SXT(−) *V. cholerae*, which should have a selective disadvantage within the clinical context since it lacks the antibiotic resistances conferred by SXT ICEs. However, SXT(−) *V. cholerae* provide a population-wide benefit, as phage that have propagated on an SXT(−) host would lack appropriate epigenetic modifications for subsequent infection of SXT ICE(+) hosts, allowing for persistence of *V. cholerae*. The prevalence of diverse anti-phage systems carried by other SXT ICEs suggests that these dynamics may be playing out in other Gammaproteobacteria.

The ability SXT ICEs to restrict beneficial mobile genetic elements such as CTXϕ and PLEs has implications on bacterial evolvability. Each SXT ICE tested restricted transduction of some, but not all, elements tested, linking the relative abundance of a given MGE or prophage to the prevalence of a restrictive SXT ICE in the population, and further demonstrating the extent to which phage predation can alter the pool of genes in a population. The ability for SXT ICEs to restrict MGEs and phages shows a tradeoff that may favor SXT(−) *V. cholerae* in certain conditions, such as in changing environments (48). SXT ICEs create a barrier to genetic exchange from cells possessing a different SXT ICE (or no SXT ICE), as only DNA modified due to activity of the same hotspot 5-encoded modification-based restriction system will be permitted as it would be indistinguishable from self, which could result in genetic divergence between closely related strains. Indeed, comparative genomics has shown that genetic exchange between genomes is higher if they encode cognate RM systems (49). Coupled with earlier findings that an SXT ICE closely related to ICE*Vch*Ind5 can limit natural transformation in *V. cholerae* through a hotspot 4 encoded DNase (50), the role of SXT ICEs in controlling bacterial gene flux is likely multifaceted.

The human gut may represent an environment where SXT ICE conjugation can occur between gut commensals or invading pathogens, especially when DNA damaging antibiotics, such as ciprofloxacin, have been administered, as ciprofloxacin has been both shown to increase SXT conjugation through activation of the SOS response (39) and is frequently secreted from cholera patients(51). Here we have shown that phage infection can increase the frequency of SXT conjugation in *V. cholerae*. Interestingly, infection by a phage that is not restricted by ICE*Vch*Ind5 stimulated higher frequency conjugation, while infection by a restricted phage did not. There is an expected energetic cost and accompanying growth delay associated with the formation of the conjugative machinery, as has been shown for other ICEs (52). When infected by a restricted phage, there may be little benefit for SXT ICEs to trigger conjugation, as their host cell may suffer a selective disadvantage limiting the SXT ICE’s vertical transmission. However, during productive phage infection, an SXT ICE that can escape a dying cell would have be able to move laterally. Together, we can now appreciate that both antibiotics and phages can spread the genes for antibiotic resistance and phage resistance by stimulating the transfer of SXT ICEs. As we continue to grapple with increasing multidrug resistant bacterial infections, which have garnered a renewed interest in phage therapy, it is of critical importance that we work to understand how phages and antibiotics interact in driving the evolution of dangerous pathogens like *V. cholerae*.

## Acknowledgments

The authors are especially thankful to icddr,b hospital and lab staff for support. The authors thank members of the Seed lab for critical feedback and thoughtful discussion regarding this manuscript and Vivek Mutalik for sharing the *E. coli* phages.

## Author contributions

K.N.L: Conceptualization, Investigation, Writing – Original draft preparation. S.G.H.: Investigation, Visualization. A.A.: Formal analysis, Software. A.C.M.: Investigation. F.T.J.: Resources, Visualization. M.S.: Resources, Project Administration. T. A: Resources, Supervision. M.A.: Resources, Supervision. K.D.S.: Conceptualization, Investigation, Writing – Review & Editing, Project administration, Funding acquisition. All authors discussed the results, commented on and approved the final manuscript.

## Data and materials availability

All data are available in the manuscript or the supplementary materials.

