## Supplementary material for "Temporal Shifts in Antibiotic Resistance Elements Govern Virus-Pathogen Conflicts": Methods and Supplementary Figures

### Materials and Methods

Strains and Growth Conditions

Stool samples were collected following institutional review with written consent from participants or for guardians of young participants. Stool samples collected from suspected cholera patients attending icddr,b Dhaka hospital, and Government Health Complex of Mathbaria, Pirojpur were screened for *V. cholerae* serogroup O1 and/or O139 using Crystal VC Rapid Dipstick test (RDT; Span Diagnostics, Surat, India). RDT-positive stool samples were de-identified, given a number, mixed with glycerol (to 20% final volume), stored at -80°C and shipped to University of California, Berkeley for further analysis. For isolation of *V. cholerae* onsite in Bangladesh, the RDT-positive stool samples were enriched in alkaline peptone water (APW) (pH-8.4, Difco, Detroit, MI) for 6-8 h at 37°C; and *V. cholerae* was isolated by culturing overnight on taurocholate tellurite gelatin agar (TTGA) (Difco). Typical *V. cholerae*-like colonies on TTGA were selected and confirmed using a combination of biochemical and serological methods, as described previously (Alam et al., 2006).

Additional isolation of *V. cholerae* and phages from stool was performed at the University of California, Berkeley. Initial isolation of *V. cholerae* was completed by streaking from stool directly on thiosulfate-citrate-bile salts-sucrose agar and Vibrio ChromoSelect agar (Sigma). Stool samples were also used to inoculate liquid outgrowth in APW at 30°C and 37°C with samples struck out from liquid cultures on thiosulfate-citrate-bile salts-sucrose agar and Vibrio ChromoSelect agar plates 6 and 24 hours after inoculation. After 24 hours, liquid outgrowths were aliquoted into cryovials with glycerol and frozen. Colonies appearing to be *V. cholerae* were colony purified at least twice on agar plates before confirmation by PCR and subsequent whole genome sequencing. These *V. cholerae* isolates were then added to the laboratory collection and used to isolate phages. For phage isolation, *V. cholerae* hosts were grown to mid-log, exposed to stool or APW outgrowths, and plated on 0.5% LB top agar. plaques were picked and plaque purified twice before confirmation by PCR and/or whole genome sequencing.

A detailed list of strains used in this study is found in Table S1. Isogenic strains of each SXT ICE were generated by conjugations into *V. cholerae* E7946 or *Escherichia coli* MG1655 backgrounds (as described below). Bacteria were routinely grown on LB agar plates and in LB broth with aeration at 37°C. Antibiotics were supplemented as appropriate at the following concentrations: 32μg/ml trimethoprim, 75 μg/ml kanamycin, 100 μg/ml spectinomycin, 100 μg/ml streptomycin, 2.5μg/ml chloramphenicol (*V. cholerae* on plates), 1.25μg/ml chloramphenicol (*V. cholerae* in liquid), 25μg/ml chloramphenicol (*E. coli*). For induction from plasmids expressing *gp21* or *orbA*/*gp25*, constructs were induced with 1mM 1 mM isopropyl β-D-1-thiogalactopyranoside (IPTG) and 1.5mM theophylline (Sigma) for 20 minutes prior to adding phage, and inducers were maintained in the agar overlays.

Phage spot plates were performed as a visual aid but not used to quantify phage titers, and were performed as described previously (McKitterick and Seed, 2018). Briefly, mid-log *V. cholerae* was added to 0.7% molten LB agar poured on a solid agar plate and allowed to solidify for ten minutes. Ten-fold dilutions of phage were applied to the surface in 3μL spots and allowed to dry. Plates were incubated at 37°C for 6-8 hours prior to visualization.

Whole genome sequencing and Genomic Analysis

*Vibrio cholerae* exconjugants and isolates from patient stool were prepped for whole genome sequencing using commercially available kits (Qiagen® DNeasy Blood and Tissue Kit and/or Monarch® Genomic DNA Purification Kit from New England BioLabs). Preparation of phage isolates was done as described previously (Mckitterick et al., 2018). WGS was performed by the QB3 Genomics Core at the University of California, Berkeley, or by the Microbial Genome Sequencing Center, at the University of Pittsburgh.

Determining phage-host range with phages directly from stool

*V. cholerae* isolates from patients stool samples (described above) were used as bacterial hosts to assay for susceptibility to ICP1 phages from stool. A small amount of frozen stool sample collected on a pipette tip was added to 50µl of APW, which was then serially diluted and plated with *V. cholerae* hosts using a soft-agar overlay method. *V. cholerae* isolates were added to 8mL of molten 0.7% LB agar and poured onto a large petri dish. The agar-bacterial host plate solidified at room temperature for 10 minutes before a dilution series of each stool sample was spotted onto the bacterial lawn. Spots were allowed to dry fully before placing into the incubator at 37°C for 6 to 8 hours. The presence/absence of plaquing in Figure 1B was determined from the two biological replicates of this assay with all hosts.

To verify the identity of the phages from patient stool samples, plaques were picked from both biological replicates of spot plates and resuspended in 50ul of STE. 10µl of the phage-STE sample was boiled for 10 minutes to release the phage DNA and use as template in a PCR verify the identity of phages. A multiplex PCR was performed with unique primer sets to differentiate ICP1, ICP2 and ICP3 (see Table S2 for primer list). Only samples with ICP1 were used in generating the time-shift assay results matrix.

Picked ICP1 plaques from stool were propagated on *V. cholerae* E7946 (which does not possess an SXT ICE). Three plaques were picked from each original passaged plaque and these phages were probed for the ability to re-infect and form plaques on the panel of *V. cholerae* isolates from patient stool samples identified above. Spot assays were performed as previously, and scored to generate the matrix in Figure 4.

Generation of Mutant Strains

Bacterial mutants were generated using splicing by overlap extension PCR of upstream and downstream regions of homology flanking a frt-spectinomycin-frt cassette. PCR products were introduced by natural transformation in *V. cholerae* as described (Dalia, Lazinski and Camilli, 2014), and the frt cassette was resolved following expression of a pMMB67EH derivative plasmid expressing the flp recombinase leaving a short frt scar in place. All mutations were verified by Sanger sequencing. Plasmids were constructed with Gibson Assembly.

Conjugations

Transfer of SXT/R391 elements from *Vibrio cholerae* into *V. cholerae* or *E. coli* was performed as described previously (Burrus & Waldor, 2003). Briefly, overnight donor cultures were grown in LB supplemented with trimethoprim and recipient strains were grown with appropriate antibiotics (either kanamycin or spectinomycin). Equal volumes (500µl) of donor and recipient cultures were added to a 1.5mL microcentrifuge tube and pelleted at 5000 x *g* for 3 minutes. Pellets were resuspended in 50µl LB and pipetted onto either an LB agar plate or a 0.8µm filter disk placed on top of an LB agar plate. Conjugations were allowed to proceed for 6 hours at 37°C, after which time the plate was flooded with 1mL of LB and gently scraped up with a 1000µl pipette and a dilution series was plated on either trimethoprim and kanamycin or trimethoprim and spectinomycin to select for recipients that acquired the SXT ICE. Mock conjugations were performed which excluded either donor or recipient to ensure that no spontaneous double antibiotic resistant colonies appeared. The presence of the entire SXT ICE in the *V. cholerae* E7946 background was further verified by whole genome sequencing of at least two exconjugants for each ICE*Vch*Ind6, ICE*Vch*Ind5 and ICE*Vch*Ban9.

Phage or mitomycin C (MMC) stimulated conjugations were performed as followed. Overnight cultures of donor *V. cholerae* (either KL316/317) grown in appropriate antibiotics were split into 4X 1mL tubes and the following treatments applied: mock infected (added appropriate volume of phage diluent), ICP1$^{2006_E}$ at a multiplicity of infection (MOI)~0.5, ICP1$^{2017_D}$ at MOI~0.5 or MMC (20ng/µl). Prior to treatment, an aliquot of the overnight donor was plated for quantification of colony forming units (CFUs) in order to back-calculate the MOI of the phage added. Following addition of treatments, all tubes were incubated at 37°C with aeration for 15 minutes. After 15 minutes, 0.5mL of the treated culture was mixed with 0.5mL of a differentially antibiotic marked recipient lacking the O1 antigen (KL322 or KL626) in a 1.5mL microcentrifuge tube. Cells were pelleted at 5000 x *g* for 3 minutes and resuspended in 50µl LB. The resuspended pellets were placed on a 0.8µm filter disk on an LB agar plate. Conjugations proceeded for 6hr, at which point the filter disk was removed from the plate with sterile tweezers and placed into 1ml LB in a 2mL microcentrifuge tube. The tube was vortexed for 30 seconds to detach cells from the filter. An aliquot of cells was plated on selective agar to quantify total number of viable donor cells at the end of the conjugation. To select for and enumerate exconjugants, cells were plated on trimethoprim with the appropriate antibiotic and incubated at 37°C overnight.

Efficiency of Plaquing Assays

The efficiency of plaquing (EOP) for each phage was calculated by comparing the number of plaques that a given phage forms on SXT ICE (-) host relative to the number of plaques formed on an otherwise isogenic SXT (+) host. Each EOP was calculated in triplicate, and the limit of detection is the point at which the phage is unable to productively infect the SXT (+) host while still forming plaques on a SXT (-) host. For these assays bacteria, *V. cholerae* or *E. coli*, were grown to mid-log, phage were added and given 10 minutes for adsorption at room temperature before the mixture was added to molten 0.5% LB agar (with the exception that T7 plaques were enumerated using 0.8% LB agar) with each dilution plated on a single petri dish.

Computational Approaches

To determine the prevalence of the HS5 contents from ICE*Vch*Ind5, ICE*Vch*Ind6 and ICE*Vch*Ban9 in taxa other than *V. cholerae*, the nucleotide sequence spanning the start of the WYL-domain to the end of the final ORF before the start of *traI* (Fig. 1D) was queried against the NCBI non-redundant nucleotide database using the BLASTn algorithm in October 2020. All hits in non-*V. cholerae* taxa with greater than 75% query coverage and 90% nucleotide identity were considered to be HS5 homologs.

Phylogenetic trees were constructed from nucleotide alignments done using the program MUSCLE of the conserved WYL-domain containing gene and the conjugative relaxase *traI*. Trees were built using PhyML with 100 bootstrap iterations, and visualized using FigTree v1.4.4.

A database of *Vibrio cholerae* genomes was constructed by downloading all assemblies from the relevant NCBI genome directory (https://www.ncbi.nlm.nih.gov/genome/browse/#!/prokaryotes/505/) and by a literature search for any isolated and sequenced *V. cholerae* genomes in published articles. These accessions were subsequently downloaded from the NCBI Sequence Read Archive (https://www.ncbi.nlm.nih.gov/sra) in fastq format and assembled with spades v3.14.0 using default settings (Bankevich et al. 2012). The database was refined to include only toxigenic strains as determined by being positive for both ctxB (accession:CP001235 region: 1646195-1646569) and the O1-antigen (accession:LT906614; region: 245143-278060) by BLASTn with ≥80% length and ≥80% identity. Representative SXT ICEs (Data S3) were then queried against every contig with ≥ 1kb length and ≥5x coverage (if coverage was known) from every genome assembly using BLASTn. The total length of each BLASTn hit was summed and the percent identities were averaged for each SXT. The SXT variant with the highest length was considered to be present in a genome if its total hit length was ≥ 80% of the SXT's sequence length and the average percent identity was ≥ 90% of the SXT sequence.

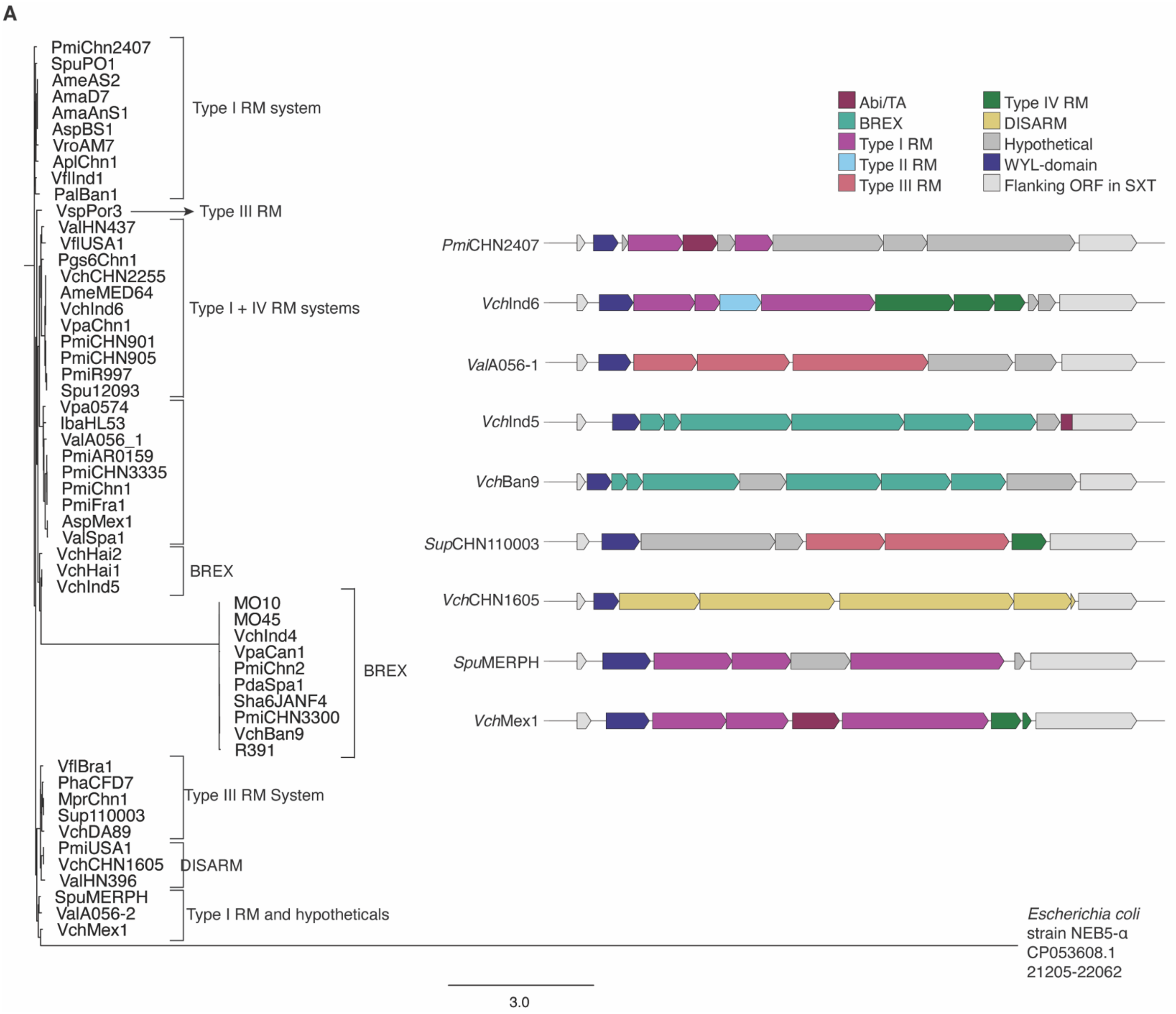
A
PmiChn2407
SpuPO1
AmeAS2
AmaD7
AmaAnS1
AspBS1
VroAM7
AplChn1
VflInd1
PalBan1
Type I RM system
VspPor3
Type III RM
ValHN437
VflUSA1
Pgs6Chn1
VchCHN2255
AmeMED64
VchInd6
VpaChn1
PmiCHN901
PmiCHN905
PmiR997
Spu12093
Type I + IV RM systems
Vpa0574
IbaHL53
ValA056_1
PmiAR0159
PmiCHN3335
PmiChn1
PmiFra1
AspMex1
ValSpa1
VchHai2
VchHai1
VchInd5
BREX
MO10
MO45
VchInd4
VpaCan1
PmiChn2
PdaSpa1
Sha6JANF4
PmiCHN3300
VchBan9
R391
BREX
VflBra1
PhaCFD7
MprChn1
Sup110003
VchDA89
Type III RM System
PmiUSA1
VchCHN1605
ValHN396
DISARM
SpuMERPH
ValA056-2
VchMex1
Type I RM and hypotheticals
Escherichia coli strain NEB5-α CP053608.1 21205-22062
3.0
Abi/TA
BREX
Type I RM
Type II RM
Type III RM
Type IV RM
DISARM
Hypothetical
WYL-domain
Flanking ORF in SXT
PmiCHN2407
VchInd6
ValA056-1
VchInd5
VchBan9
SupCHN110003
VchCHN1605
SpuMERPH
VchMex1

B

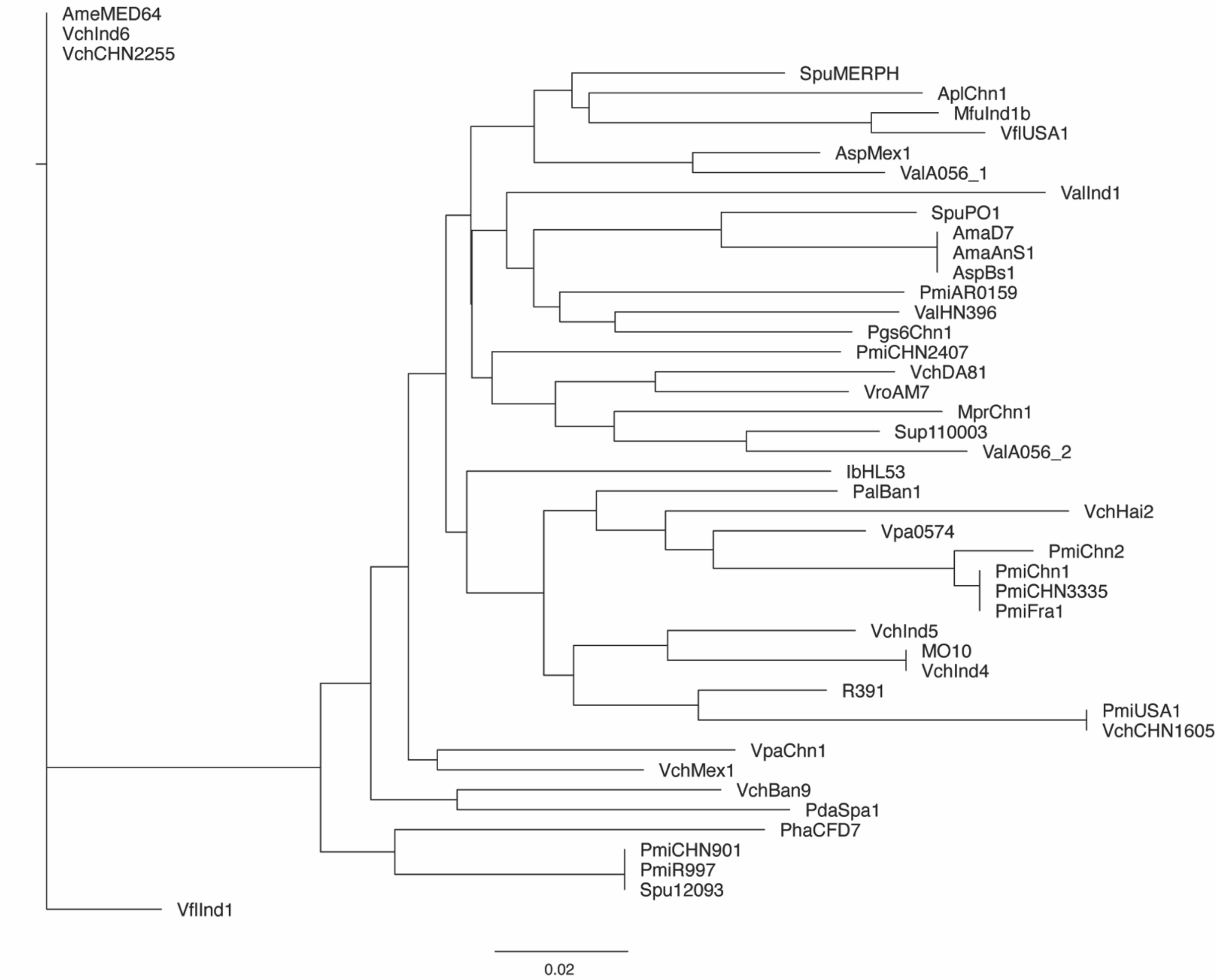


**Fig. S1. Hotspot 5 in SXT ICEs encode a conserved WYL-domain protein**. (A) Neighbor-joining phylogenetic tree from a MUSCLE alignment of the nucleotide sequence for the WYL-domain containing gene, showing SXT ICE groups based on predicted anti-phage systems encoded by hotspot 5, with the WYL-domain protein from *Escherichia coli* indicated as an outgroup. Gene maps for each representative node are depicted showing the upstream hypothetical s025 and downstream conserved *traI* genes as flanking genes in hotspot 5. (B) Neighbor-joining phylogenetic tree from a MUSCLE alignment of the nucleotide sequence for conserved downstream gene *traI* showing no relationship between hotspot 5 anti-phage type system and similarity.

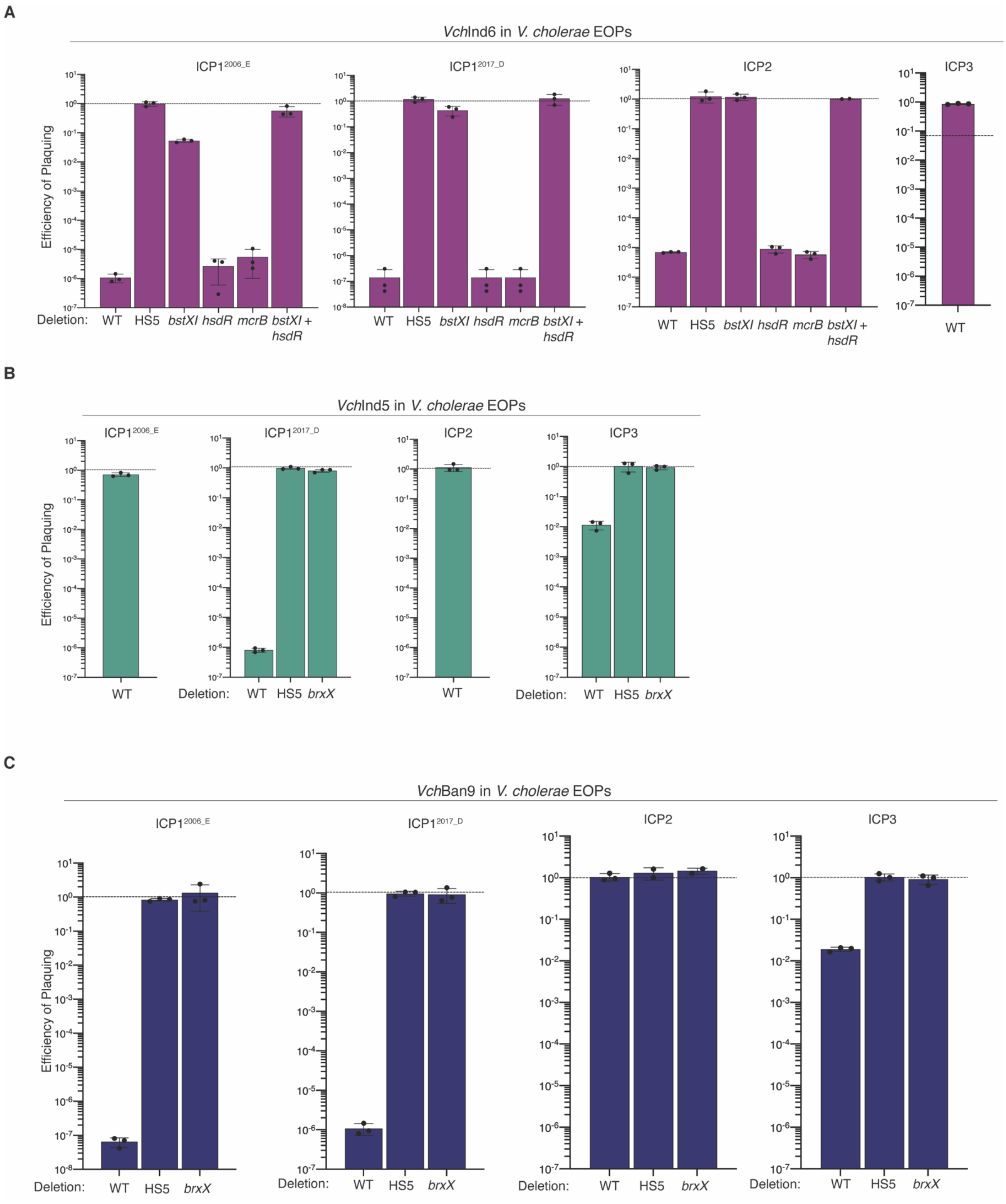
A
VchInd6 in V. cholerae EOPs
ICP1 2006_E
ICP1 2017_D
ICP2
ICP3
Efficiency of Plaquing
Deletion: WT HS5 bstXI hsdR mcrB bstXI + hsdR
WT
B
VchInd5 in V. cholerae EOPs
ICP1 2006_E
ICP1 2017_D
ICP2
ICP3
Efficiency of Plaquing
WT
Deletion: WT HS5 brxX
C
VchBan9 in V. cholerae EOPs
ICP1 2006_E
ICP1 2017_D
ICP2
ICP3
Efficiency of Plaquing
Deletion: WT HS5 brxX

**Fig. S2. SXT ICEs confer protection against lytic vibriophages.** Source data for Figure 2B in which wild type and SXT ICE knockouts were conjugated into *V. cholerae* E7946 and the efficiency of plaquing (EOP) was calculated for the lytic phages $ICP1^{2006\_E}$, $ICP1^{2017\_D}$, ICP2, ICP3 as the average of triplicate plaque assays. The dotted line indicates an EOP of 1, where plaquing is the same for a permissive *V. cholerae* E7946 SXT(-) host as the SXT(+) exconjugant. (A) EOPs for phages on ICE*Vch*Ind6 *V. cholerae*. (B): EOPs for phages on ICE*Vch*Ind5 *V. cholerae*. (C) EOPs for phages on ICE*Vch*Ban9 *V. cholerae*.

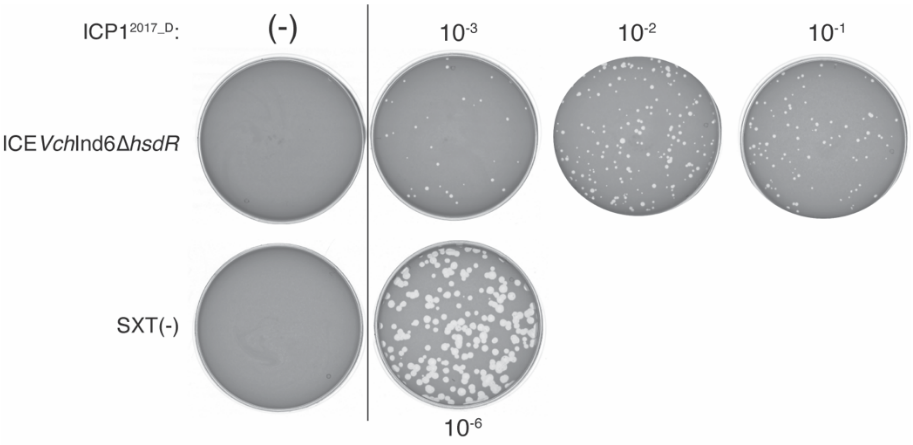


**Fig. S3. ICE*Vch*Ind6 *hsdR* is necessary for full inhibition of ICP1$^{2017_D}$.** Plaque assays of ICP1$^{2017_D}$ on *V. cholerae* ICE*Vch*Ind6Δ*hsdR* (top) and SXT(-) (bottom). Uninfected, far left. Dilution series of ICP1$^{2017_D}$ from $10^{-3}$ up to $10^{-1}$ (top) and $10^{-6}$ on SXT(-). Results are representative of assays performed in triplicate.

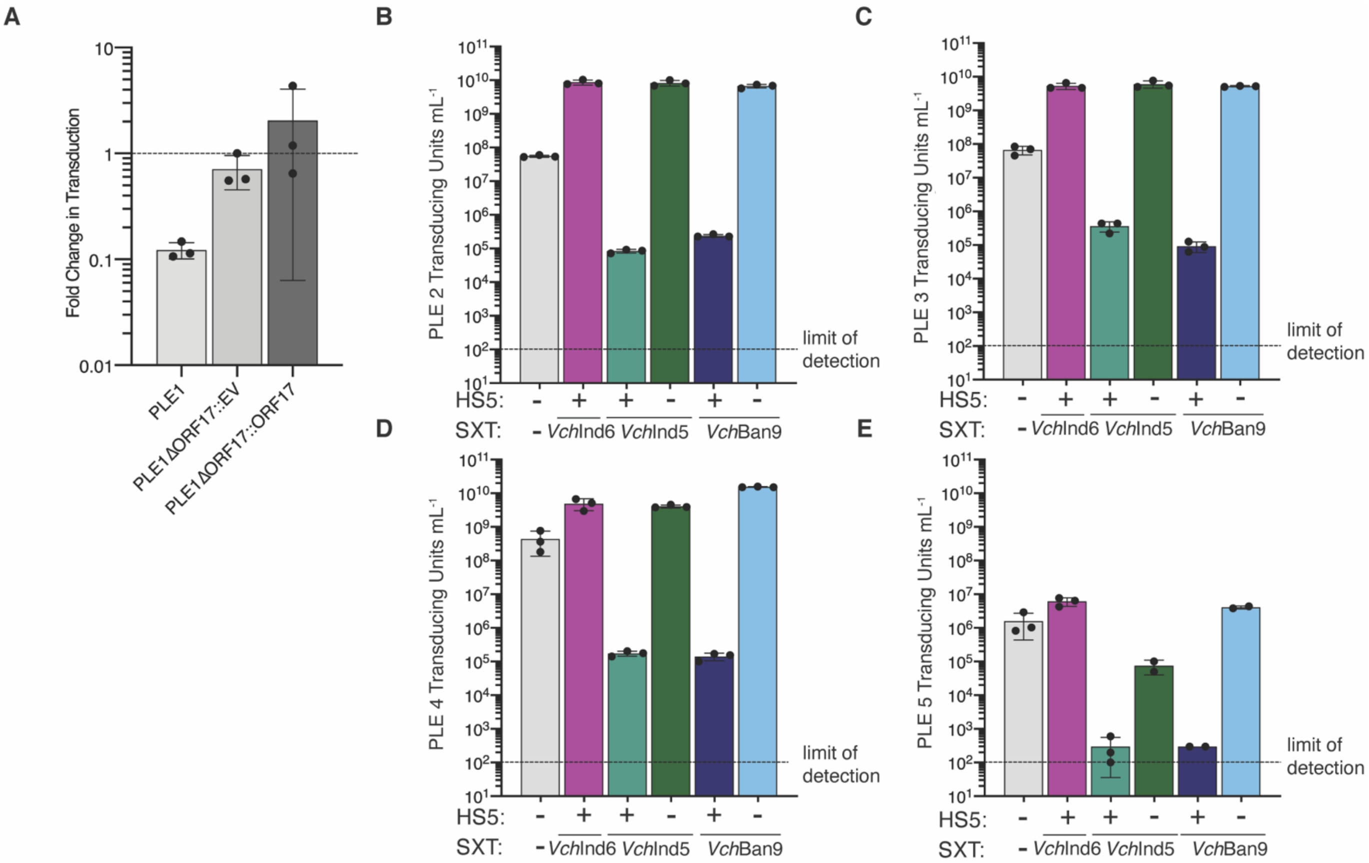


**Fig. S4. SXT ICEs limit acquisition of PLEs: mobile anti-phage elements** (A) Transduction of wild type PLE 1 or PLE1Δ*orf17* containing the putative recognition motif for ICE*Vch*Ind6-encoded BstXI. PLE 1Δ*orf17* was complemented with either an empty vector or *orf17 in trans*. Transductions were done into either into an SXT(-) or ICE*Vch*Ind6(+) host. (B-E): Transductions of PLEs 2-5 into *V. cholerae* E7946 harboring SXT ICEs. Both wild type and hotspot 5 deletions were assayed as recipients. ICE*Vch*Ind6 did not inhibit PLEs 2-5 so the ICEVchInd6ΔHS5 strain was not tested.

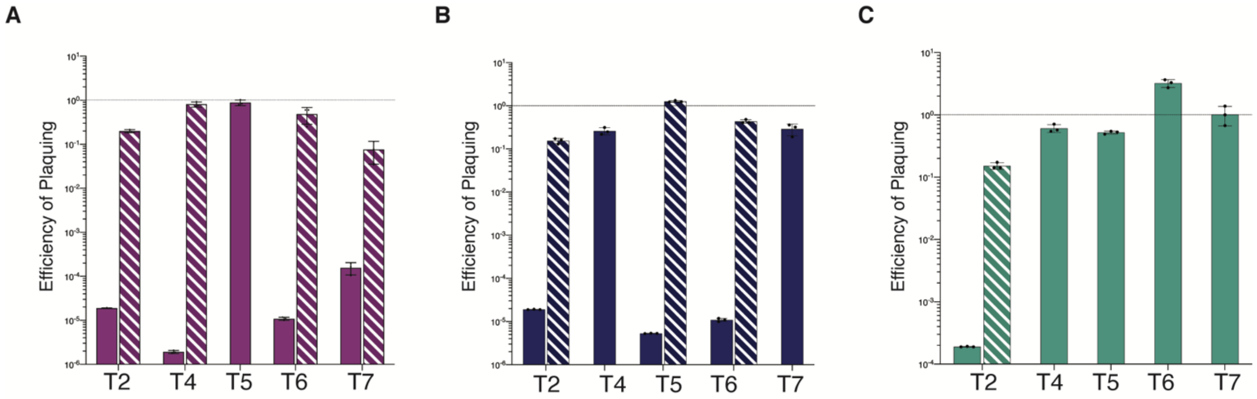


**Fig. S5. SXT ICEs confer protection against lytic phage in *E. coli*.** Source data for Figure 3. Wild type SXT ICEs (solid bars) and hotspot 5 (HS5) knockouts (patterned bars) were conjugated into *E. coli* MG1655 and the efficiency of plaquing (EOP) was calculated for the lytic phages T2, T4, T5, T6 and T7 as the average of triplicate plaque assays. The dotted line indicates and EOP of 1, where plaquing is the same for a permissive SXT (-) host as the SXT (+) exconjugant. (A) ICE*Vch*Ind6 (solid bars) and ICE*Vch*Ind5ΔHS5 (patterned lines). (B) ICE*Vch*Ind5 (solid bars) and ICE*Vch*Ind5ΔHS5 (patterned lines) (C) ICE*Vch*Ban9 (solid bars) and ICE*Vch*Ban9ΔHS5 (patterned lines).

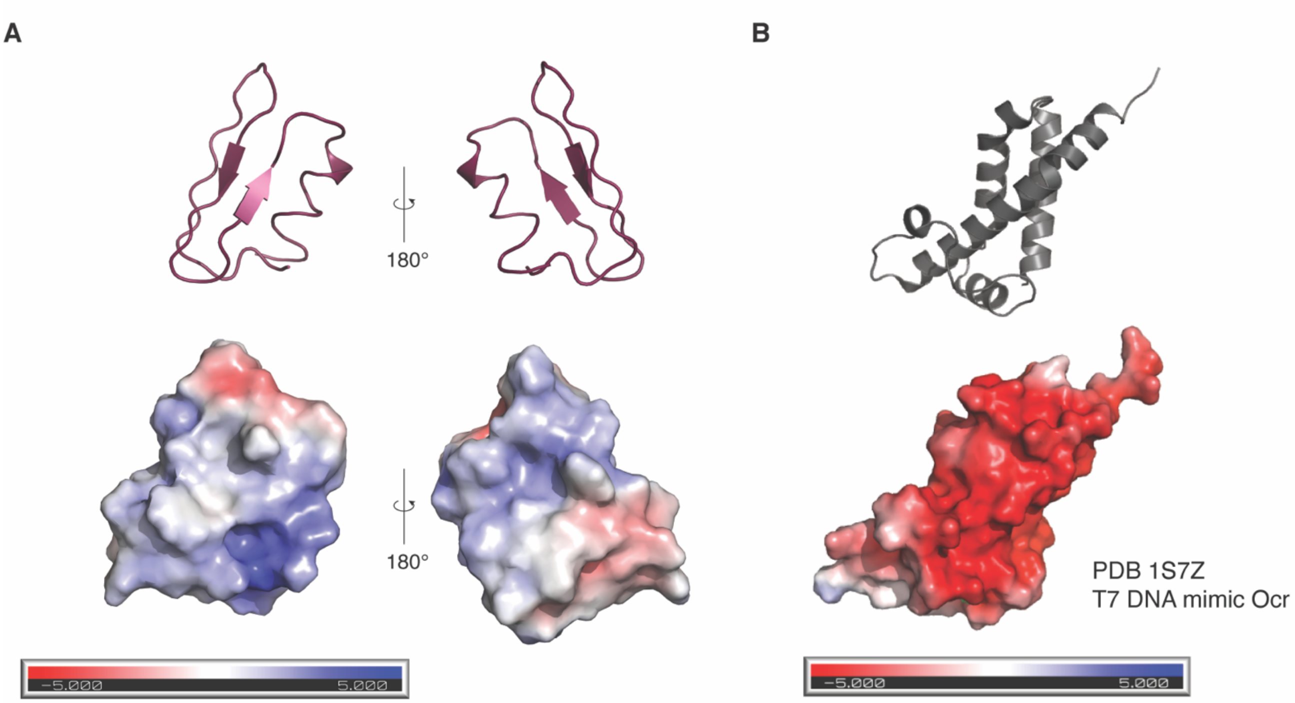


**Fig. S6. ICP1 encoded OrbA predicted structure and surface electrostatic potential map**. (A) Predicted structure of OrbA from i-TASSER, showing 180 degree rotation of around the vertical axis showing predicted beta sheets and predicted helix, with APBS electrostatics overlayed indicating negative charged surfaces with red and positively charged surfaces in blue. (B) Solved crystal structure of the known anti-BREX protein T7 DNA mimic Ocr (PDB 1S7Z) (Walkinshaw et al., 2002) with electrostatics overlayed on the structure.

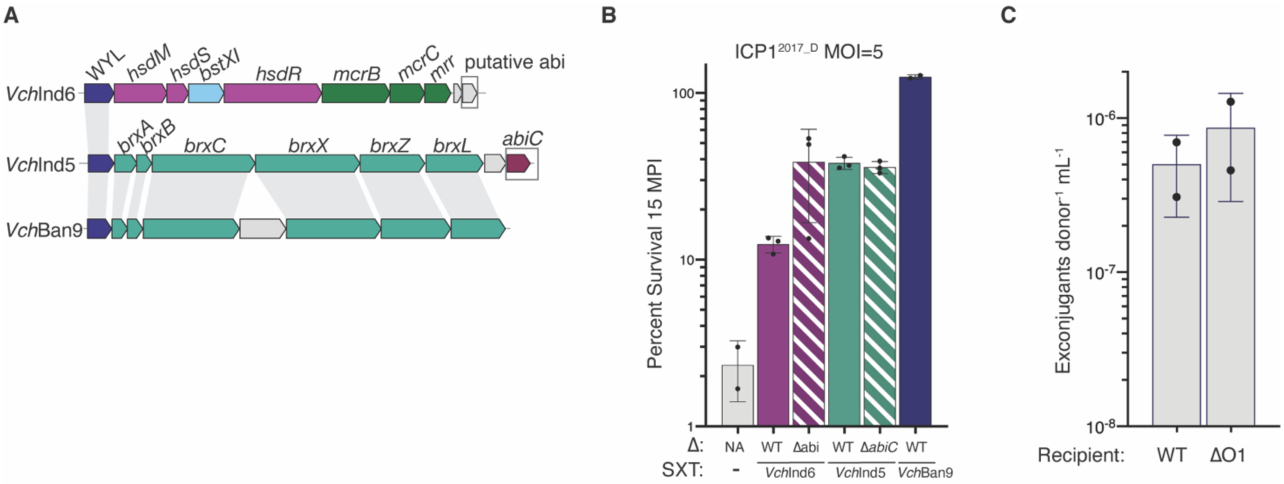


**Fig. S7. Impact of phage infection on an isogenic exconjugants of SXT ICEs** (A) Genomic organization of hotspot 5 in SXT ICEs tested showing the genes encoding putative abortive infection proteins in grey boxes. (B) Cell survival 15 minutes after an MOI=5 infection by ICP1$^{2017_D}$ of *V. cholerae* E7946 SXT(-), or derivatives harboring ICE*Vch*Ind6, ICE*Vch*Ind5, ICE*Vch*Ban9 and single gene knockouts of putative abortive infection genes indicated.

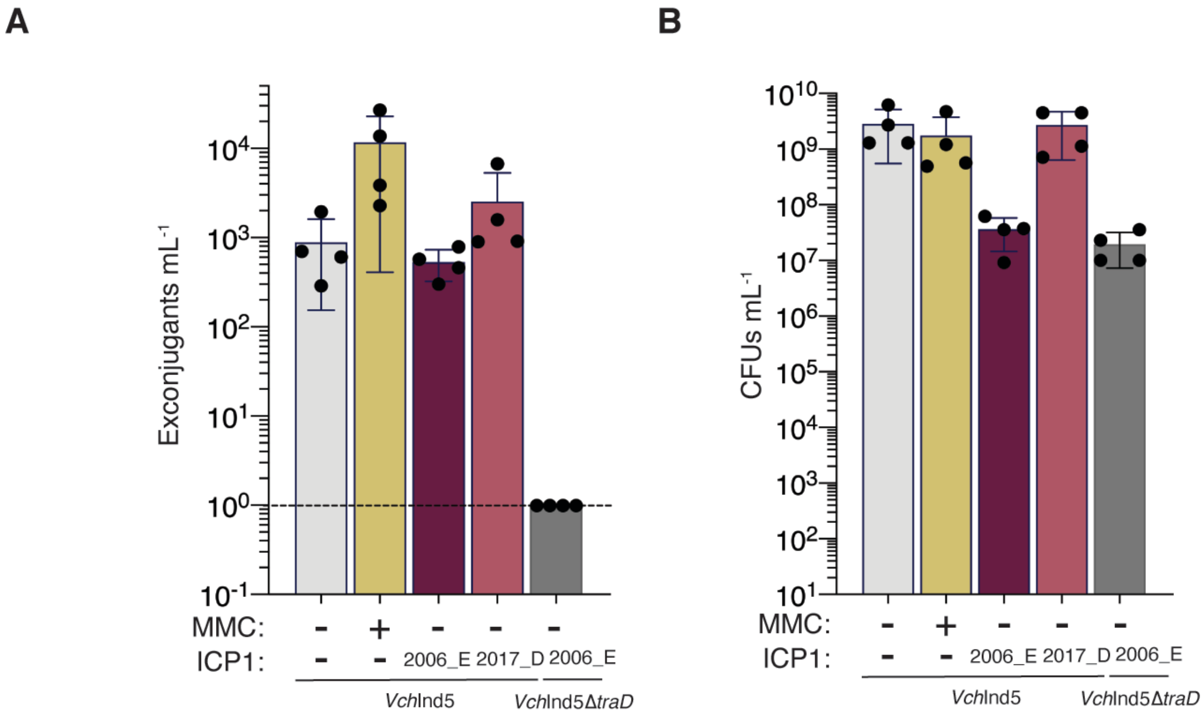


**Fig. S8. ICP1 stimulated SXT ICE conjugation results in cell death but equivalent levels of conjugation.** (A) Raw number of exconjugants obtained following conjugation with donor ICE*Vch*Ind5(+) *V. cholerae* treated with mitomycin C (MMC), or infected by ICP1$^{2006_E}$ and ICP1$^{2017_D}$ prior to conjugation compared to an untreated control. (B) Enumeration of surviving ICE*Vch*Ind5(+) *V. cholerae* donors treated with mitomycin C (MMC), or infected by ICP1$^{2006_E}$ and ICP1$^{2017_D}$ following conjugation compared to an untreated control.
